# Conserved phosphorylatable residues in motif G of positive-stranded virus RdRps regulate polymerase activity and suggest targets for drug design

**DOI:** 10.64898/2025.12.08.692995

**Authors:** Camille Duflos, Belén Lizcano-Perret, Olve B. Peersen, Gaëtan Herinckx, Yeranddy A. Alpizar, Didier Vertommen, Thomas Michiels

**Affiliations:** Université catholique de Louvain, de Duve Institute, Brussels, Belgium; Department of Biochemistry & Molecular Biology, Colorado State University, Fort Collins, CO 80523-1870, USA; MASSPROT platform, de Duve Institute, Université Catholique de Louvain, Brussels, Belgium; KU Leuven Department of Microbiology, Immunology & Transplantation, Rega Institute, Division of Virology, Antiviral Drug & Vaccine Research, Laboratory of Molecular Vaccinology & Vaccine Discovery (MVVD); Leuven, Belgium

**Keywords:** polymerase, RdRp, phosphorylation, virus, Theiler’s murine encephalomyelitis virus, TMEV, severe acute respiratory syndrome, SARS-CoV-2, positive-stranded RNA virus, drug target

## Abstract

RNA viruses rely on an RNA-dependent RNA polymerase (RdRp) to replicate their genome. RdRps share a conserved core replicase structure described as a right hand within which RNA replication occurs. RNA polymerases contain a series of conserved motifs (A-G) that are essential for catalysis. Motif G, located at the RNA entry channel, guides the incoming RNA into the catalytic centre and and holds it in place during catalysis. Although RdRp phosphorylation has been reported, it has been scarcely studied. In most studied cases, phosphomimetic mutations reduced viral replication.

In this study, we identified Theiler’s murine encephalomyelitis virus (TMEV) polymerase (3D^pol^) residues that undergo some extent of phosphorylation in infected cells. Among these residues, Thr109 and Ser110 located in motif G are highly conserved in the sequences of picornavirus polymerases and in the structure of many positive-stranded virus polymerases, including nsp12 of severe acute respiratory syndrome coronavirus 2 (SARS-CoV-2). Using mutagenesis and reporter viruses, we show that phosphomimetic mutation of either residue abrogates viral replication, for both TMEV and SARS-CoV-2. Mutations of 3D^pol^ residues 109 and 110 into all other possible residues shows that, besides negatively charged phosphomimetic residues, bulky residues strongly inhibit replication, suggesting that phosphorylation inhibits polymerase activity by steric hindrance and/or through charge repulsion with RNA entering the catalytic core. Because these phosphorylatable residues are surface-exposed and conserved among viral polymerases, they represent promising targets for the rational design of broad-spectrum antiviral agents.

**Importance:** RNA viruses require an RNA-dependent RNA polymerase to replicate their genome. We identified in Theiler’s murine encephalomyelitis virus polymerase (TMEV 3D^pol^) residues that undergo some extent of phosphorylation in infected cells. Among these residues, Thr109 and Ser110 are located in the entry channel of the polymerase, a region important for directing the RNA into the polymerase and locking it in place during catalysis. Incidentally, Thr109 and Ser110 are highly conserved in the sequences of picornavirus polymerases and in the structures of many positive-stranded virus polymerases including severe acute respiratory syndrome coronavirus 2 (SARS-CoV-2) nsp12. Our study revealed that, in both TMEV 3D^pol^ and SARS-CoV-2 nsp12, mutation of either residue into a negatively charged amino acid that mimics phosphorylation abrogates viral replication, suggesting phosphorylation would block polymerase activity. As these phosphorylated residues are accessible and conserved, they provide important candidate targets for the design of antiviral molecules.

## Introduction

RNA viruses rely on their own machinery to replicate and transcribe their genomes within host cells. Central to the viral replication machinery are the RNA-dependent RNA polymerases (RdRp), which share a common catalytic activity embedded in a conserved right hand shaped core architecture [1]. RdRps can also have functions other than genome replication carried out by additional modules that are part of the polymerase itself, by a larger partially processed polyprotein or by interacting co-factors. Polymerase activity can also be regulated by post-translational modifications such as phosphorylation that may modulate activity, stability, subcellular localization, and interactions with host or viral proteins [2, 3].

Central to the activity of the polymerase are the hallmark conserved motifs A to G. Their shape and/or key residues ensure the functions of the polymerase. The template RNA enters the polymerase from the “top” and contacts motifs G and F that contain key residues which guide the incoming RNA into the active site and lock it in place during the nucleotide addition cycle. Motif F also forms the ceiling of the NTP entry tunnel. Once inside, the incoming NTP is held in place by motifs A and B while motifs A and C coordinate the two metal ions necessary for catalysis (Fig. 1) [4, 5].

**Figure 1.**
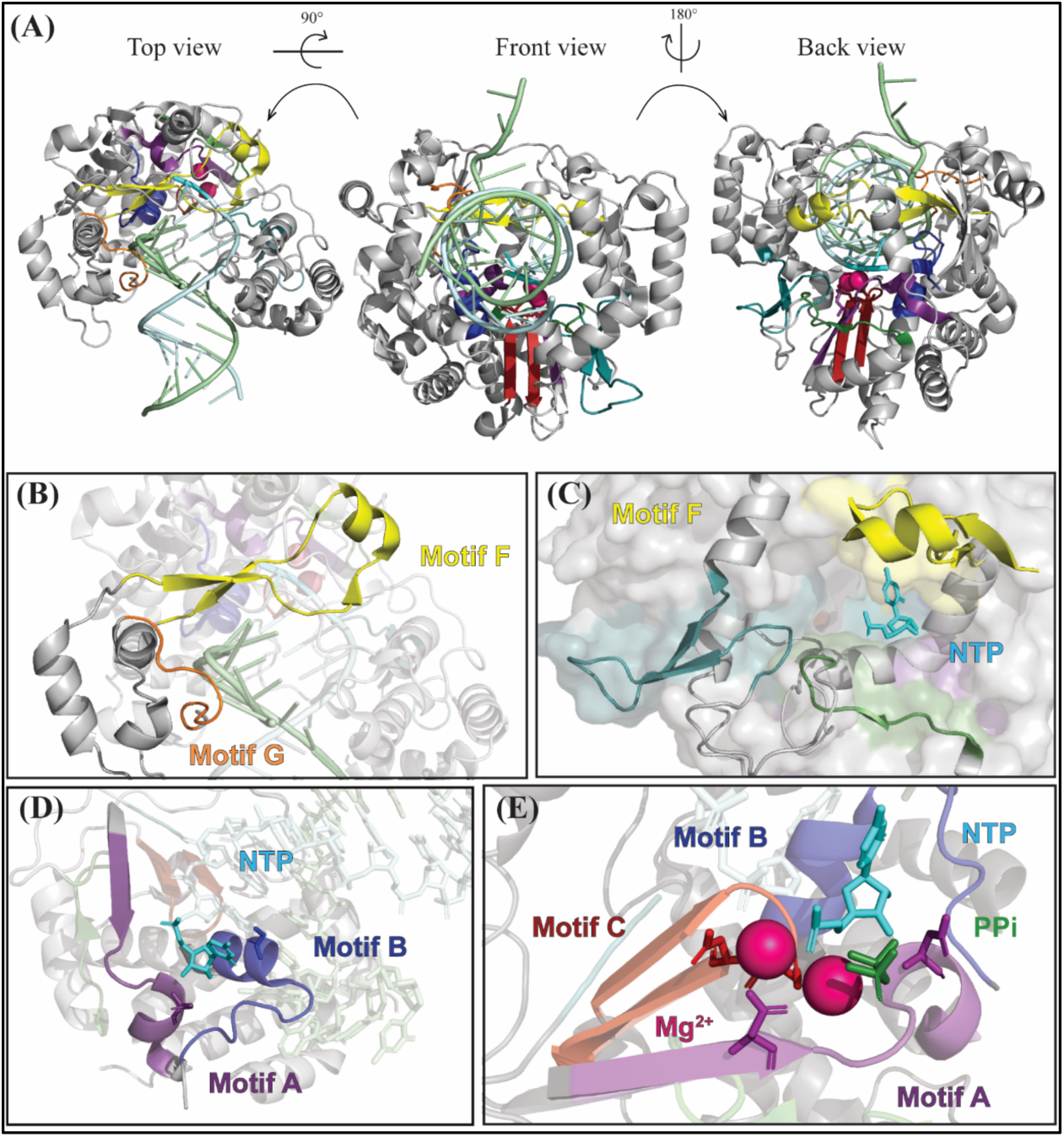
Structure of a typical picornavirus polymerase based on poliovirus (PV) 3D^pol^ (PDB: 3OL7 [10]) **A.** PV 3D^pol^ seen as a cartoon from the “top”, “front” and “back”. The different motifs, metal ions and RNA are coloured: motif A (purple), B (blue), C (red), D (green), E (teal), F (yellow), G (orange); Mg^2+^ (magenta), template RNA (pale green), product strand (pale cyan), newly incorporated nucleotide (cyan) and pyrophosphate (green). **B.** PV 3D^pol^ “top” view, zoomed on the entry canal. Motifs F and G and the template RNA are put forward. **C.** Surface representation of 3D^pol^ seen from the back, motifs D, E and F (cartoon) are put forward showing the NTP entry channel. The newly incorporated NTP is in sticks representation. **D.** PV 3D ^pol^ “top” view, zoomed on the catalytic core. Fingers, thumb and Mg^+2^ ions were removed for clarity. Motifs A and B are put forward, especially the residues important for the positioning of the incoming NTP (sticks). **E.** PV 3D^pol^ “top” view, zoomed on the catalytic core. Fingers and thumb were removed for clarity. The newly incorporated NTP and pyrophosphate as wells as motifs A and C are put forward, especially the residues that coordinate the Mg^2+^ ions (sticks).

Theiler’s murine encephalomyelitis virus (TMEV) and severe acute respiratory syndrome coronavirus-2 (SARS-CoV-2), represent distinct positive-stranded RNA viruses with contrasting genome organizations and replication strategies.

TMEV is a member of the genus *Cardiovirus* within the family *Picornaviridae* and has an 8 kb-long RNA genome which, upon entry in the cell cytosol, gets translated by cellular ribosomes into a polyprotein thanks to an internal ribosome entry site (IRES). This polyprotein is mainly processed by viral protease 3C^pro^ thus releasing the various viral proteins including 3D^pol^ the RdRp. Persistent TMEV strains such as the DA strain used in this work have a striking ability to infect the murine central nervous system and can persist despite a strong and specific immune response. The virus can infect neurons but is mostly detected in oligodendrocytes and macrophages in chronically infected mice, the natural host of TMEV [6].

SARS-CoV-2 is an enveloped virus with a 30kb-long genome. It primarily infects the human respiratory tract, although infection has also been detected in patients’ heart, central nervous system and other organs, especially in severely ill patients [7–9]. The coronavirus genome is a capped, polyadenylated RNA containing several open reading frames (ORFs). Upon delivery into the cytosol, ORF1 can be translated as a polyprotein, the processing of which yields non-structural proteins nsp1 to nsp11 (ORF1a) or proteins nsp1 to nsp16 after a ribosomal frame shift (ORF 1ab). These proteins are mainly implicated in genome replication, with nsp12 being the polymerase. After genome replication, transcription by nsp12 gives rise to several sub-genomic RNAs (sgRNAs) allowing the expression of structural proteins such as the spike, membrane, envelope and nucleocapsid (S, M, E, N) as well as other less characterized proteins.

Starting from the observation that Thr109 and Ser110 residues, located in TMEV’s 3D^pol^ motif G, can be phosphorylated during infection and that these phosphorylatable residues are conserved in the polymerases of many positive-stranded RNA viruses, we explored the effects of phosphorylation on TMEV 3D^pol^ and SARS-CoV nsp12.

## Results

### TMEV 3D^pol^ can be phosphorylated in infected cells

Phosphorylation of TMEV 3D polymerase (3D^pol^) was unexpectedly detected by mass spectrometry in infected HeLa M cells during an unrelated study. We sought to confirm the presence of this phosphorylation in cells from the natural host of this virus by infecting murine fibroblast derived L929 cells with virus KJ6, a variant of the TMEV DA strain carrying a capsid adapted to optimize L929 cell infection [11]. In the chronic phase of TMEV infection, macrophages become the most abundant virus reservoir although the infection is somehow limited in these cells [6]. We thus also infected macrophage derived RAW264.7 cells with virus VV18, a macrophage adapted strain [12]. Cells were infected for different periods of time after which crude lysates or immunoprecipitated 3D^pol^ were in-gel trypsin digested and analyzed by mass spectrometry in order to identify phosphorylated residues. We observed the phosphorylation of the following 3D^pol^ residues at least once in this series of experiment: Ser10, Ser63, Thr109 or Ser110, Ser389 and Ser406 (**Table 1**). Since Thr109 and Ser110 are adjacent residues, it was not possible to determine with certainty which of them was phosphorylated.

**Table 1.**
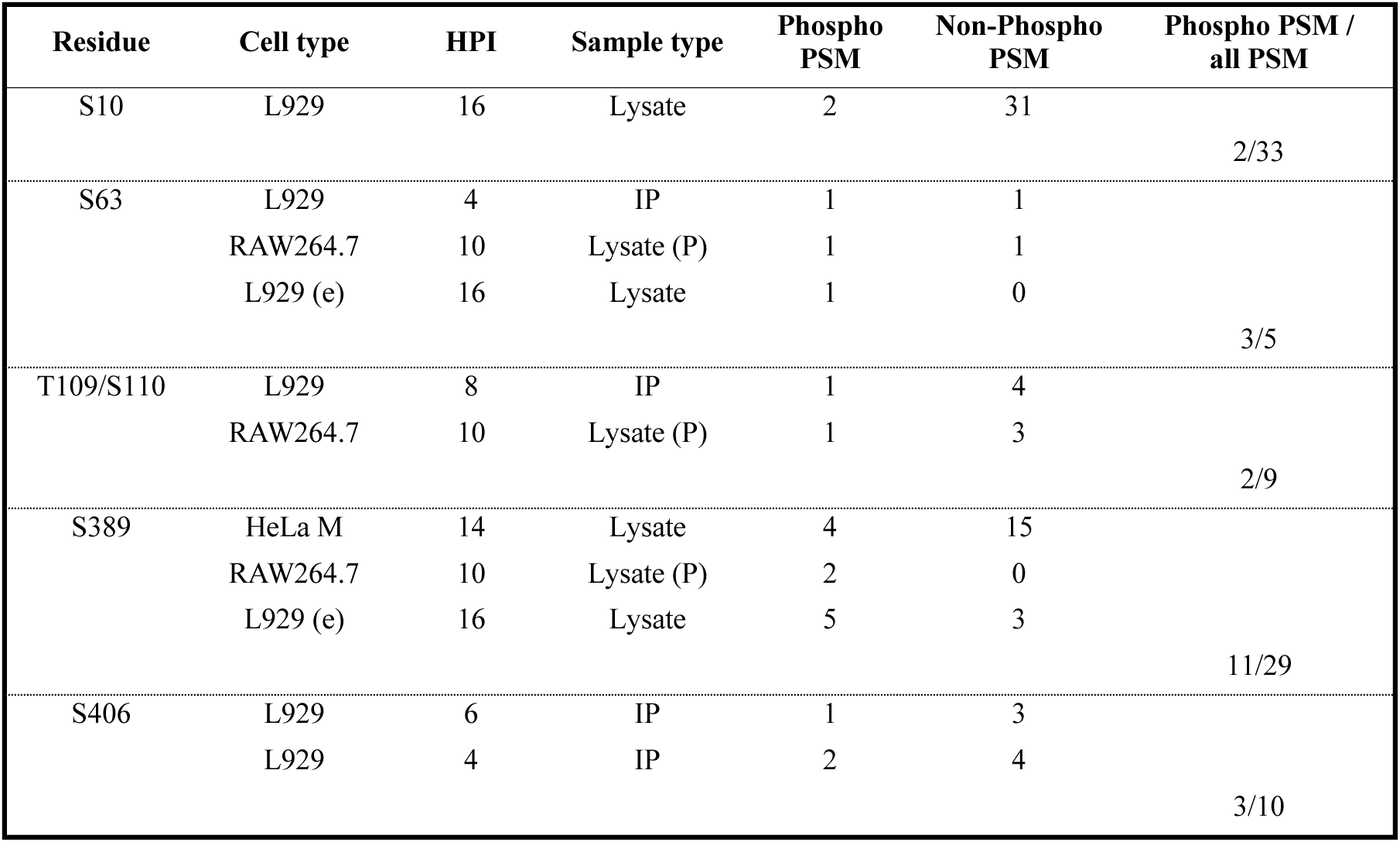
TMEV 3D^pol^ phosphorylated residues detected by mass spectrometry analysis. Cells were infected with WT TMEV (VV18 variant at 10 PFU per cell for RAW264.7 cells and KJ6 variant at 2 PFU per cell for L929 or HeLa M cells) for variable times of infection. Cells were lysed, either in Laemmli buffer ("lysate") or in RIPA buffer for subsequent immunoprecipitation (IP) of 3D^pol^. Some RIPA lysate pellets (lysate P) were also analyzed. Crude lysates, IP products or RIPA lysis pellets were migrated on SDS gels, in gel trypsin digested and submitted for mass spectrometry analysis. HPI= hours post infection, (e) = epoxomycine treatment, PSM = peptide spectrum match.

| Residue | Cell type | HPI | Sample type | Phospho PSM | Non-Phospho PSM | Phospho PSM / all PSM |
| --- | --- | --- | --- | --- | --- | --- |
| S10 | L929 | 16 | Lysate | 2 | 31 | 2/33 |
| S63 | L929 | 4 | IP | 1 | 1 | 3/5 |
|  | RAW264.7 | 10 | Lysate (P) | 1 | 1 |  |
|  | L929 (e) | 16 | Lysate | 1 | 0 |  |
| T109/S110 | L929 | 8 | IP | 1 | 4 | 2/9 |
|  | RAW264.7 | 10 | Lysate (P) | 1 | 3 |  |
| S389 | HeLa M | 14 | Lysate | 4 | 15 | 11/29 |
|  | RAW264.7 | 10 | Lysate (P) | 2 | 0 |  |
|  | L929 (e) | 16 | Lysate | 5 | 3 |  |
| S406 | L929 | 6 | IP | 1 | 3 | 3/10 |
|  | L929 | 4 | IP | 2 | 4 |  |

Among the phospho-residues identified, only Ser406, Thr109 and Ser110 are conserved within cardioviruses 3D^pol^ (Fig. 2A). Interestingly, residues Ser110 and to a lesser extent Thr109 but not Ser406 are well-conserved across the *Picornaviridae* family (Fig. 2B).

**Figure 2.**
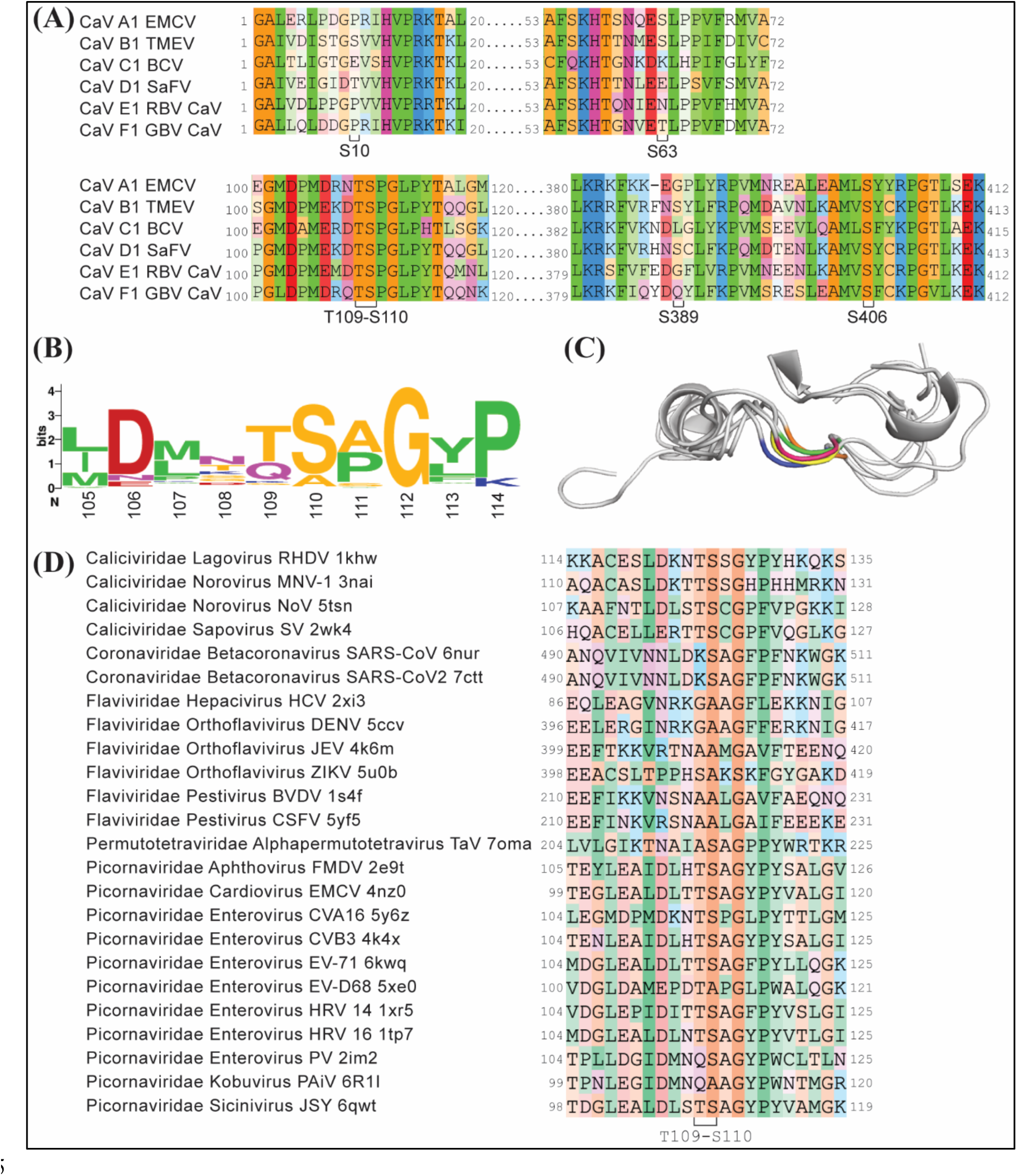
Sequence alignment of phosphorylated residues. **A.** Sequence alignment of representative cardiovirus (CaV) 3Dpol. Only residues surrounding the identified phosphorylated residues are shown. **B**. Consensus (WebLogo) of representative Picornaviridae 3Dpol sequences surrounding Thr109 Ser110. Numbering of the horizontal axis corresponds to TMEV 3Dpol residues. **C**. Cartoon representation of the structure-based sequence alignment shown in D. One polymerase of each viral family is depicted with Thr109 and Ser110 coloured: NoV (5TSN) is green, SARS-CoV-2 (7CTT) is orange, DENV (5CCV) is yellow, TaV (7OMA) is pink and EMCV (4NZ0) is blue) **D**. Structure-based sequence alignment of the (+)ssRNA RdRps found in the PDB based on [1]. Ten amino acids before and after Thr109-Ser110 are shown. Only structures where motif G were crystalized and aligned with 4NZ0 were included in the analysis. Residue numbering corresponds to that used in the PDB structures. EMCV= Encephalomyocarditis virus; BCV= Boone Cardiovirus; SaFV= Saffold virus; RBV= Red-backed vole cardiovirus; GBV= Grey-backed vole cardiovirus; RHDV= Rabbit hemorrhagic disease virus; MNV= Murine Norovirus; NoV= Norwalk virus; SV= Sapporo virus; HCV= Hepatitis C virus; DENV= Dengue virus; JEV= Japanese encephalitis virus; ZIKV= Zika virus; BVDV= Bovine viral diarrhea virus; CSFV= Classical swine fever virus; TaV= Thosea asigna virus; FMDV= Foot-and-mouth disease virus; CV (A16-B3)= Coxsackievirus A16 and B3; EV (71-D68)= Enterovirus 71 or D68; HRV= Human rhinovirus; PV= Poliovirus; PAiV= porcine Aichi virus; JSY= Sicinivirus strain JSY

Moreover, a structural alignment performed on the RdRp structure dataset published by Peersen [1, 13] (version: v4), showed a fair conservation of these residues amongst polymerases of positive-stranded RNA viruses. Residues matching Thr109 are phosphorylatable threonine in several viruses and residues matching Ser110 are always serine or alanine, which is found mainly in the *Flaviviridae* family (Fig. 2C and D). These residues are located in polymerase motif G at the entrance of the template RNA tunnel (Fig. 1A and B)[4].

### Phosphomimetic mutations of TMEV 3D^pol^ Thr109 and Ser110 inhibit TMEV replication

The conservation of residues Thr109-Ser110 and their particular location in the polymerases prompted us to study the impact of the phosphorylation of Thr109 and Ser110 on TMEV 3D^pol^ activity. We first sought to produce TMEV viruses harbouring either phosphoinhibiting (Ala) or phosphomimetic (Asp or Glu) mutations at those positions by electroporation of *in vitro* transcribed viral genomic RNA into BHK-21 cells. The supernatant recovered 48-to-72h post electroporation (hpe) was titrated by plaque assay on BHK-21 cells (Fig. 3A). Viruses containing a non-phosphorylatable Ala residue at positions Thr109 or Ser110 had titers close to those of the wild type virus (8x10^4^ PFU/mL and 1x10^5^ PFU/mL, respectively compared to 5x10^5^ PFU/mL for the WT). In contrast, viruses carrying the phosphomimetic Asp or Glu mutations yielded very low titers (between 4x10^3^ PFU/mL and 7x10^3^ PFU/mL). Sanger sequencing of RT-PCR amplified viral genome segments indicated that the few viruses recovered after transfection of the phosphomimetic mutant viral RNAs corresponded to revertant viruses where the phosphomimetic residues had reverted to wild type or mutated to other amino acids (Fig. 3B), in contrast to alanine mutants, which kept the introduced Ala codon.

**Figure 3.**
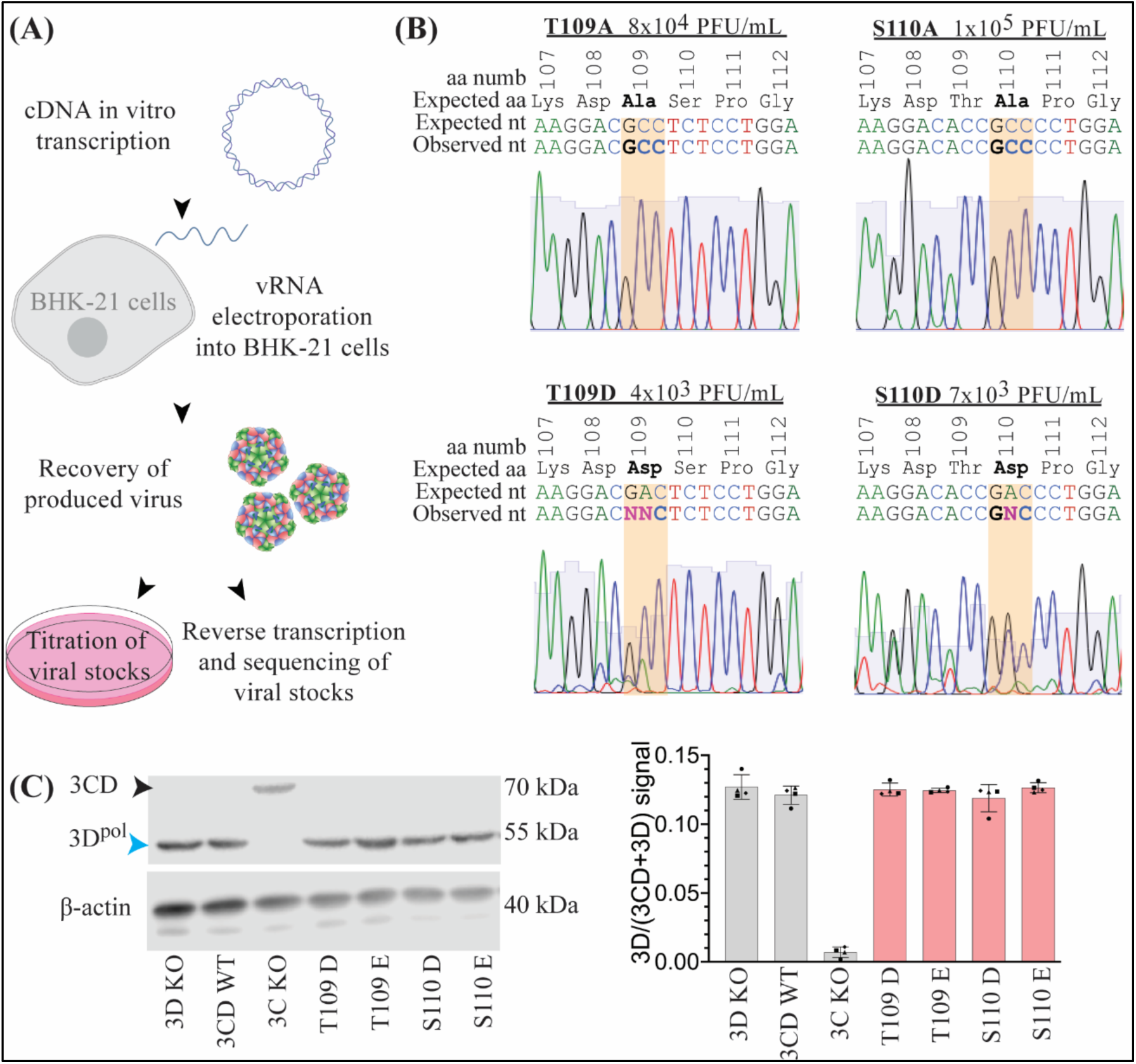
Titration and sequencing of Thr109 and Ser110 Ala and Asp mutants. **A.** Workflow of the experiment. Viral cDNA was *in vitro* transcribed and electroporated into BHK-21 cells. The recovered supernatants were titrated by plaque assay, and RNA was isolated for reverse-transcription and cDNA sequencing. **B.** Sanger sequencing of the viral stocks obtained from the different mutant viruses as well as the titers obtained for each of them. Top left panel Thr109 Ala, top right Ser110 Ala, bottom left Thr109 Asp, bottom right Ser110 Asp. Residue numbers are shown on top, with the expected residue identity followed by the expected nucleotide sequence and the one obtained. **C.** Western blot example showing 3D^pol^ or 3CD (and β-actin as loading control) detection in 293T cells, 24 h after transfection of pcDNA3 plasmids encoding for WT and mutant 3CD proteins. The right panel shows the quantification of the signal for 3 independent experiments, 3D^pol^ signal was normalized by the sum of the signals 3D^pol^ and 3CD 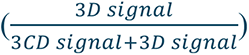

During polyprotein processing, 3D^pol^ is first embeded in its precursor form 3CD, which has protease but no replicase activity. Processing of the 3CD precursor into the mature 3C^pro^ and 3D^pol^ is done in trans by 3C^pro^ or by 3CD itself, which might be affected by our mutations.

Western blot analysis of 3CD proteins ectopically expressed in 293T cells suggested that the phosphomimetic mutations introduced in 3D^pol^ affected neither protein production nor 3C^pro^- mediated 3CD processing and thus likely affected polymerase function directly (Fig. 3C).

Taken together, our results indicate that the phosphomimetic mutations of TMEV 3D^pol^ at either position Thr109 or Ser110 impede viral replication.

The occurrence of revertants in primary viral stocks impeded proper analysis of mutant replication. To circumvent this problem, we assessed primary viral RNA replication using a split nanoluciferase assay in cells transfected with *in vitro* transcribed viral genomic RNA (vRNA) [14, 15](Fig. 4 A-B). Therefore, vRNA of viruses expressing the short nLuc fragment (HiBiT) fused to the N-terminus of the L protein was electroporated into BHK-21 cells stably expressing the large nLuc (BHK-21-LgBiT). A reporter virus with a catalytically inactive 3D^pol^ (3D^pol^ KO, motif C YGDD→YGAA mutation) served as a negative control. In a preliminary kinetics experiment, nLuc signals from replication competent and 3D^pol^ KO constructs paralleled up to 4h post vRNA electroporation (hpe) and thus reflected HiBiT production by early IRES-mediated translation of transfected vRNA and processing of the viral polyprotein (Fig. 4B). From 4 hpe nLuc signals of WT and 3D^pol^ KO viruses started diverging, showing the impact of replication on nLuc activity. Therefore, replication ability of 3D^pol^ mutants was measured as the ratio between nLuc activity measured at 18 and 4 hpe, 18 hpe reflecting replication and at 4 hpe reflecting vRNA transfection efficiency (Fig. 4A).

**Figure 4.**
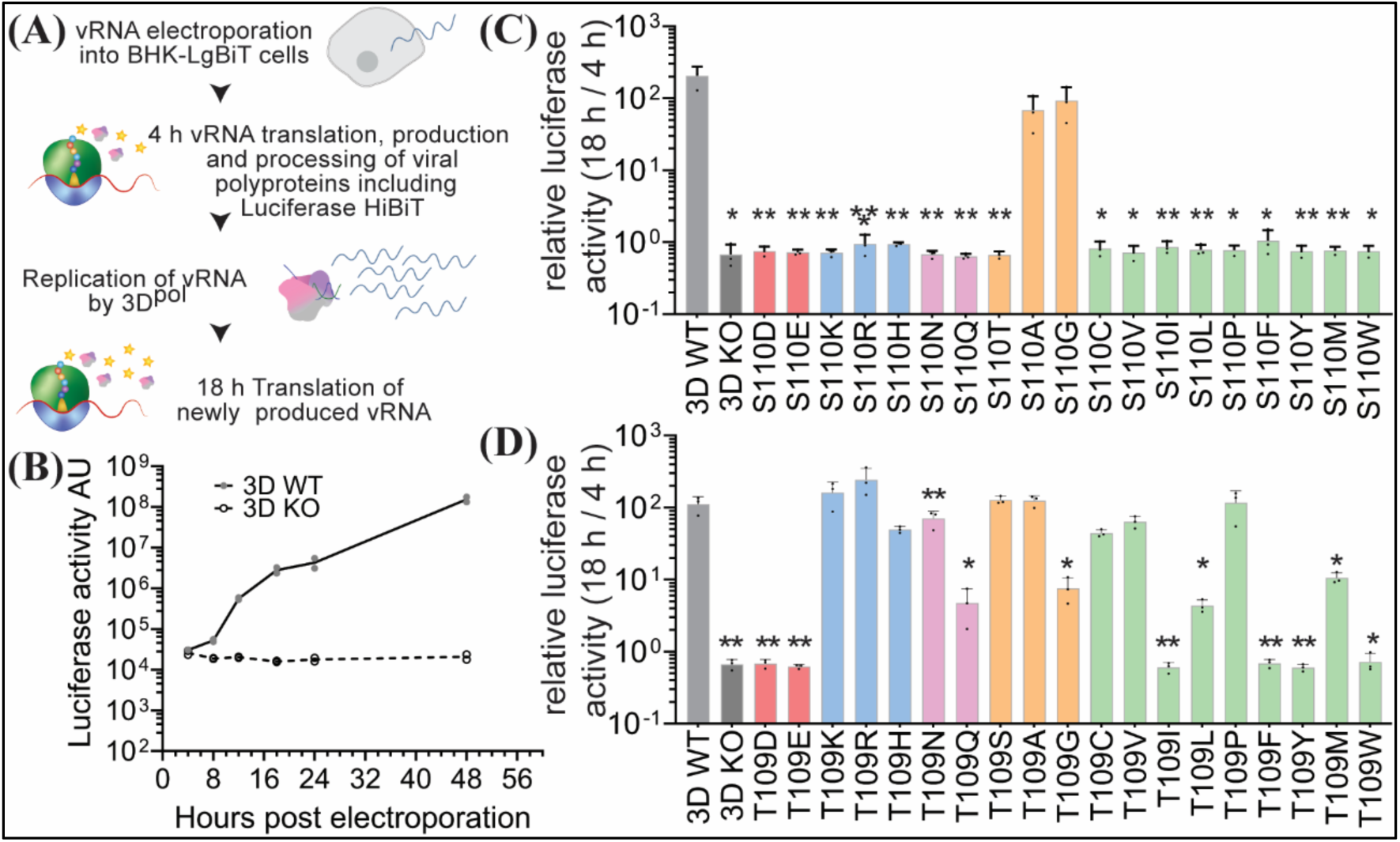
Split luciferase-mediated assessment of T109 and S110 mutant virus replication. **A.** Schematic split-luciferase viral RNA replication assay. **B.** TMEV replication time course experiment. Luciferase signals for WT and 3D^pol^ KO viruses were measured 4, 8, 18, 24 and 48 hpe. Results are shown for 2 technical replicates. Curves indicate the average of both measurements. **C and D** TMEV replication of Ser110 (C) and Thr109 (D) mutant viruses. Luciferase signal was measured after initial translation (4 hpe), and after viral replication (18 hpe).The graphs show the ratio between the luciferase activity measured at 18 and 4 hpe, each showing the average for 3 technical replicates. Bars represent the mean +/- SEM. Statistical analysis is ANOVA all *vs* WT. *=p<0.1 **p<0.01 ***p<0.001, n=3.

To define whether polymerase activity suppression by Thr109-Asp and Ser110-Asp mutations was due to the phospho-mimicking negative charge of the Asp residue, we generated a series of mutants where 3D^pol^ Thr109 and Ser110 codons were mutated into all the other possible residues (Fig. 4C and D).

### TMEV 3D^pol^ Ser110 phosphorylation likely abrogates viral replication by sterically occupying the template entry channel

Mutation of 3D^pol^ Ser110 into small residues such as Ala or Gly allowed near WT replication levels, which was not the case for the other residues (Fig. 4C). These results are in line with those of Wang et al. [16] who showed that EV71 3D^pol^ could accommodate corresponding Ser115-Ala and Ser115-Gly but not Ser115-Thr mutations.

As these residues are located in motif G at the entrance of the template RNA channel, replacement of Ser by a larger residue likely hinders the proper binding of the RNA template into the catalytic channel. Accordingly, addition of a phosphate to the Ser110 residue would likely block polymerase activity through steric hindrance.

### TMEV 3D^pol^ Thr109 phosphorylation likely abrogates viral replication by electrostatic repulsion of the template vRNA or by modifying the structure of motif G

Residue Thr109 is less stringent in terms of mutation tolerance (Fig. 4D). In line with our previous results, viruses harbouring an acidic residue at position 109 (Asp or Glu) did not show evidence of replication, with a nLuc 4/18 hpe activity ratio similar to that of the 3D^pol^ KO negative control (Fig. 4D-red bars). Asn and Gln mutants showed that in absence of net charge, small size residues are preferred over larger ones (Fig. 4D-pink bars). This conclusion can be extended to more apolar residues such as tryptophan, phenylalanine or tyrosine that do not allow for any polymerase activity while smaller ones such as valine, leucine or methionine do (Fig. 4D-green bars). The Ala mutant replicated well whereas the Gly mutant had moderate replication ability, suggesting that factors other than size are at play (Fig. 4D- orange bars). Also, Arg, Lys and His mutants replicated to or near to wild type levels, suggesting that their electropositive nature that is compatible with RNA contact might compensate for their larger size (Fig. 4D-blue bars).

To confirm the identification of residues at position 109 that support 3D^pol^ activity, we isolated and sequenced revertants occurring upon electroporation of vRNAs coding for mutant 3D^pol^ bearing phosphomimic Asp or Glu residues at position 109 and their uncharged counterparts Asn and Gln (supplemental Table A1). From the Gln mutant, which replicated to intermediate levels, most clones kept the mutation and few revertant viruses were obtained. From the Asp and Glu mutants, all recovered viruses had reversions. Residues occurring in the revertants were in good agreement with the residues found to support 3D^pol^ activity in the luciferase test. Of note, there was a bias toward reverted codons carrying only one nucleotide difference compared to the electroporated vRNA, which also explains that no reversion to WT was obtained.

Two revertants caught our attention because they had a second site Glu106-Gly mutation while retaining the Glu or Gln mutation introduced at position 109 three residues away. To gain an insight into the effect of this additional mutation, we generated 3D^pol^ models using AlphaFold (Fig. 5A). These models suggest that mutation of Thr109 into a bulky (especially negative) amino acid creates a steric clash with Glu106, which can be avoided with the mutation of Glu106 into a small residue as Gly. In the luciferase replication assay, the Glu106-Gly mutation alone did not substantially affect viral replication (Fig. 5B-grey lined bar). However, this mutation was able to mitigate the effect of dampening mutations such as Thr109-Gln, Thr109-Glu and Thr109-Leu (Fig. 5B-red, pink and green lined bars). Of note, a negative residue is almost systematically observed at a position equivalent to Glu106, in polymerases of the *Picornaviridae* family and of the closely related *Caliciviridae* family (Fig. 2D).

**Figure 5.**
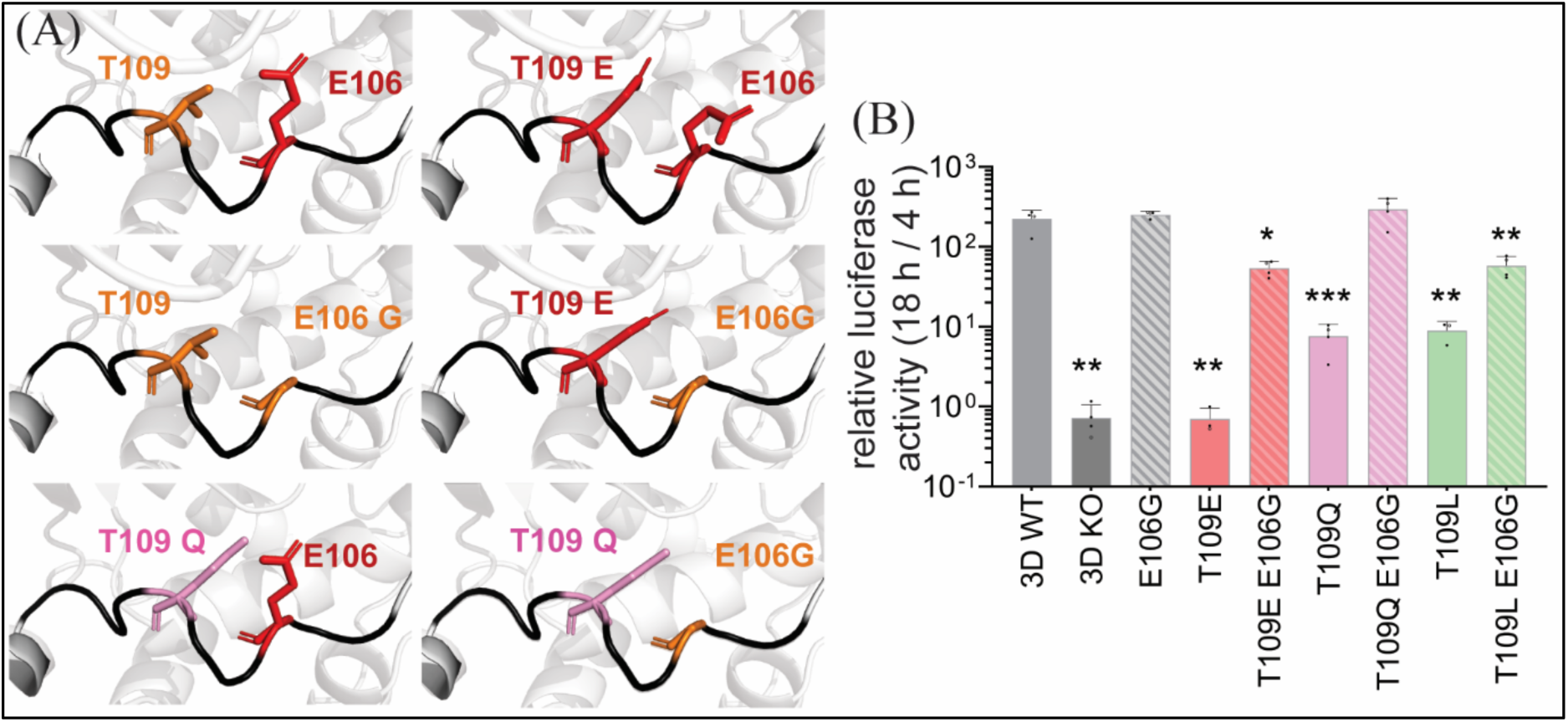
Influence of the Glu106 second-site mutation on viral replication. **A.** AlphaFold predictions of TMEV 3D^pol^ Thr109 and Glu106 mutants. Top left panel WT 3D^pol^, top right Thr109 Glu mutant, middle left panel Glu106-Gly mutant, middle right panel Thr109-Glu – Glu106-Gly double mutant, bottom left panel Thr109-Gln mutant, bottom right panel Thr109-Gln – Glu106-Gly double mutant. **B.** TMEV replication of Thr109-Glu106 mutant viruses. Graphs show the mean +/- SEM of the 18/4 hpe nLuc activity ratio measured in 3 independent experiments, each showing the average for 3 technical replicates. Statistical analysis is ANOVA all *vs* WT. *=p<0.1 **p<0.01 ***p<0.001, n=3.

Taken together, the data indicate that Thr109 and Glu106 residues play an important role in shaping motif G loop or the RNA entry canal. Our results indicate that the structure is maintained by the size and charge of both residues.

### Mutation of Thr109 into a positive residue (Arg) does not provide the virus a fitness advantage

We noticed that the Thr109-Arg mutant reproducibly, generated slightly more luciferase activity than the WT control (Fig. 4D). Strikingly, in SARS-CoV-1, SARS-CoV-2 nsp12, and the polymerase of most *Nidovirales* sequences, the residue structurally corresponding to Thr109 is a lysine residue (Fig. 2D). To study whether a positively charged residue at that position would provide a fitness advantage in specific conditions, we performed competition assays by successive passage of a 50/50 mixture of WT and mutant Thr109 Arg TMEV in L929 fibroblast, Neuro2A neuroblast and RAW264.7 macrophage murine cell lines, as well as in type I interferon-deficient hamster BHK-21 cells (Fig. 6).

**Figure 6.**
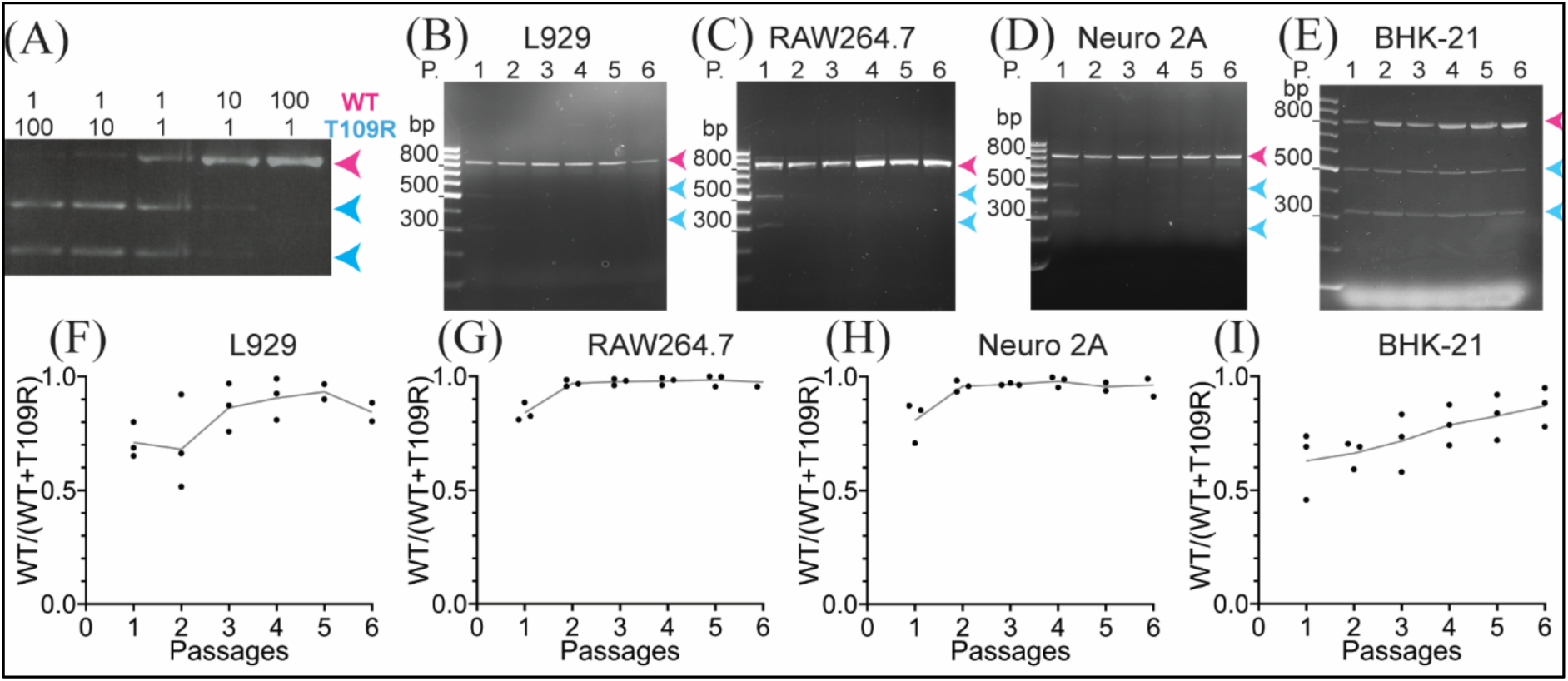
Competition between Thr109-Arg 3D^pol^ mutants and WT. **A.** T109R and WT viruses were mixed at different ratios based on their titer. The RNA from the mixes was extracted, submitted to RT-PCR and digested with BglII, thus producing a single 804 nt-long fragment for viruses with a WT polymerase (pink arrow), and 489 nt and 315 nt products for the Thr109 Arg mutant viruses (light blue arrows). The proportion of each virus is indicated above the gel. **B-I.** Cells were infected with a 1:1 virus mixture of WT TMEV and 3D^pol^ Thr109 Arg TMEV. Supernatants were recovered after appearace of cytopathic effect (CPE) and used to infect fresh cells (passage 1, P.1), the procedure was repeated for 6 passages. RNA extracted from the supernatants was subjected to RT-PCR and BglII digestion as in (A). B, to E show agarose gel examples. F, to I show the quantification results of 3 independent experiments 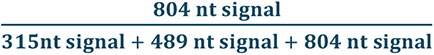. The lines connect the average of the 3 experiments. Passages were carried out in L929 (B&F), RAW264.7 (C&G), Neuro 2A (D&H) or BHK-21 (E&I) cells.

The virus carrying a positively charged mutation was consistently out-competed by the WT virus within the first two passages (Fig.6 B-D and F-H), except in BHK-21 cells where mutant virus could still be detected after passage 6 (Fig. 6E & I).

In conclusion, although replacement of Thr109 by a negatively charged phosphomimetic residue abrogates replication, replacement of this residue by a positively charged residue is tolerated but at a significant viral fitness cost.

### The effect of SARS-CoV-2 nsp12 mutations in motif G mirror those of TMEV 3D^pol^

Residues Lys501 and Ser502 of SARS-CoV-2 nsp12 structurally align with TMEV 3D^pol^ Thr109 and Ser110. We thus tested the influence of mutating those residues on the replication of a SARS-CoV-2 luciferase reporter replicon. The bacterial artificial chromosome (BAC) carrying the replicon was obtained by deleting most of the spike coding sequence in BAC pCCI-4K-SARS-CoV-2-NLuc [17], which carries an infectious clone of SARS-CoV-2 (Wuhan strain) coding for the nLuc connected to the C-terminus of the ORF7a, through a foot-and-mouth disease virus (FMDV) 2A “ribosomal skipping” linker [18]. In this construct, the cDNA of SARS-CoV-2 is preceded by a cytomegalovirus (CMV) promoter and followed by an hepatitis delta virus (HDV) ribozyme and a Simian Virus 40 terminator (SV40) [19] (Fig. 7A).

**Figure 7.**
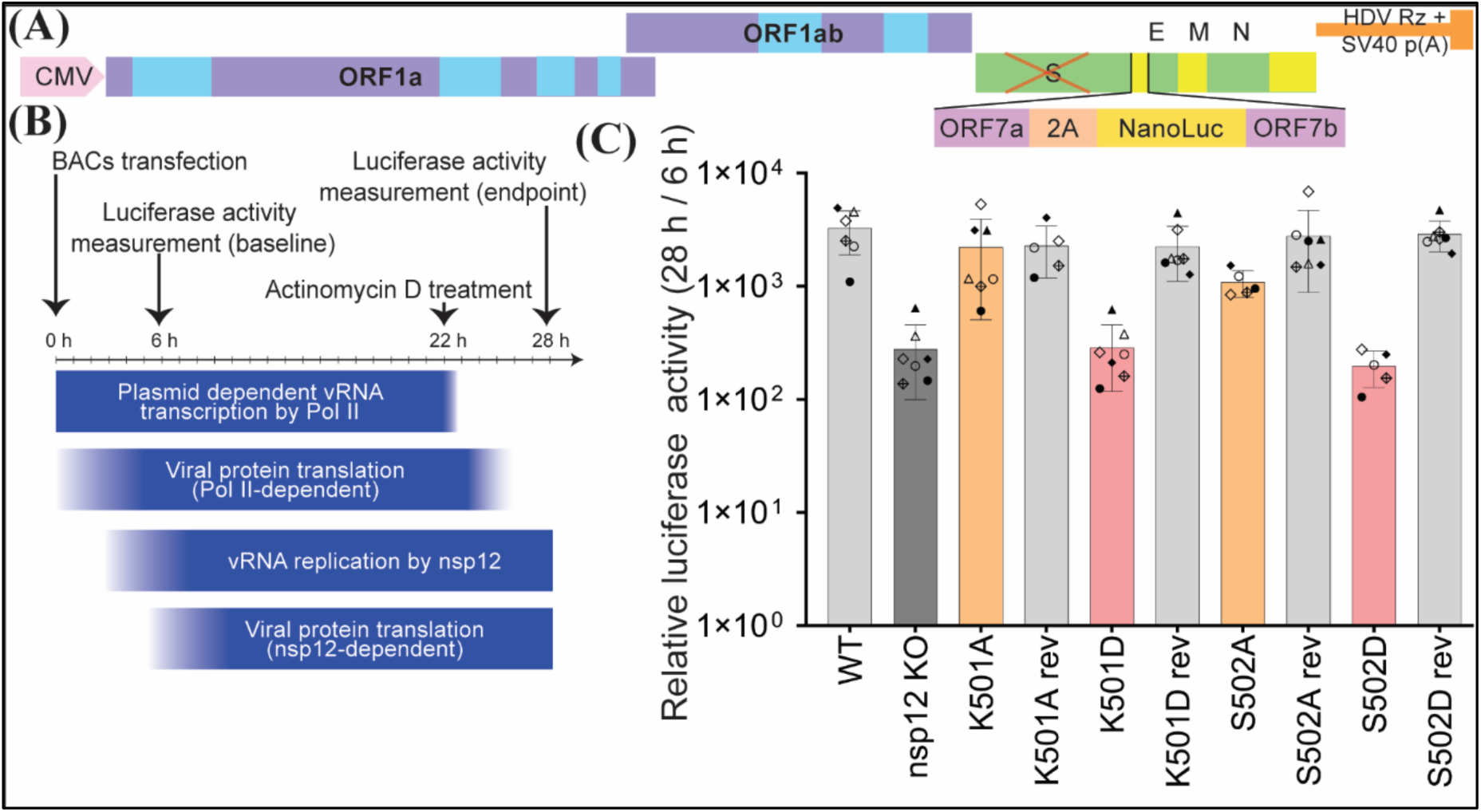
SARS-CoV-2 Luciferase assay. **A.** Schematic representation of the luciferase reporter SARS-CoV-2 replicon contained in the BAC vector. Figure adapted from [17]. **B.** Replication assay: 293T cells were transfected with the different BACs containing SARS-CoV-2 replicon cDNA. Luciferase activity was measured 6 hpt, after an initial round of Pol II-mediated transcription and of translation. 22 hpt actinomycin D was added to limit Pol II-mediated transcription and luciferase activity was measured at 28 hpt. **C.** Replication of SARS-CoV-2 mutants. n=3, each dot represents the mean ratio between the luciferase activity measured at 28 and 6 hpt, for 4 technical replicates. Each shape (diamond, circle, triangle) represents replicates done with different cDNA batches. Bars represent the average +/- SEM. Statistical analysis is ANOVA all *vs* WT. ***p<0.001, ****p<0.0001

In the BAC construct expressing the SARS-CoV-2 replicon, nsp12 mutations to Ala or Asp of Lys501 and Ser502 were introduced to assess the influence of these residues in replication. A catalytically inactive nsp12 mutant (motif C: SDD to SNN) was also produced, as a control. For each mutant generated, a revertant BAC clone was made, where the mutated region was replaced back by a wild type sequence, to ensure that observed phenotypes of the mutants are not due to mutations occurring elsewhere in the BAC constructs.

Upon transfection of this vector into 293T cells, some luciferase signal was detected even for the catalytically inactive mutant, possibly due to aberrant nsp7-2A-nLuc transcription from a cryptic promoter. Thus, nLuc activity was measured at 6 h post transfection (hpt) to control for transfection efficiency and background effects, as well as at 28 hpt, a time where replication of the SARS-CoV-2 RNA replicon increased nLuc expression (Fig. 7B). A low dose of actinomycin D was added 22 hpt to decrease the aberrant nsp7-2A-nLuc transcription by RNA Pol II once viral replication was established. Results shown in Fig. 7 are in line with those obtained for TMEV 3D^pol^, in that mutation of nsp12 residues Lys501 and Ser502 to Asp, but not to Ala, abolished nsp12 activity (Fig. 7C).

We next examined whether Ser502 of nsp12 could be phosphorylated in infected cells by producing a rabbit polyclonal anti-nsp12 antibody to enable immunoprecipitation from infected cells. A549 cells expressing hACE2 and TMPRSS2 were infected with SARS-CoV-2 Wuhan strain at an MOI of 2 and nsp12 was immunoprecipitated from the cell lysates 48 hours after infection, in-gel trypsin digested, and submitted to mass spectrometry analysis. In two independent replicates, nsp12 peptides were detected via 29 (38% coverage) and 21 (29% coverage) different peptides. One peptide spectrum match containing Ser502 was detected in each experiment, but its phosphorylated counterpart could not be observed and thus our study did not allow us to confirm the phosphorylation status of nsp12.

## Discussion

In this work we observed that TMEV 3D^pol^ is phosphorylated at several sites, one of them being the conserved motif G, a flexible loop located at the RNA template entry channel of the polymerase. Although motif G is described as being mainly structural with little sequence conservation [4], structure-based sequence alignments showed that Ser110 of TMEV 3D^pol^ is conserved amongst many +ssRNA viral polymerases while adjacent Thr109 is less conserved, being most often a small or positive residue (Fig. 2D).

Interestingly, in EV-71 3D^pol^-RNA complexed crystals, Ser115 (the homolog of TMEV 3D^pol^ Ser110) was proposed to lock the template in position for catalysis by creating a steric blockage in the backbone of template nucleotides n+1 and n+2. It is only by overcoming this blockage that the template can translocate and allow for the following round of nucleotide addition cycle [16]. Given the structural similarity between all picornavirus polymerases, it can reasonably be proposed that RdRp residues Ser110 (TMEV) and Ser115 (EV71) play the same role, as supported by our extensive mutagenesis results (Fig. 4). As suggested by the lack of replication of viruses carrying phosphomimetic mutations, phosphorylation of Ser110 in infected cells most likely inhibits viral replication.

The role of Thr109 in polymerase function is harder to interpret. Amino acids with a net negative charge (Asp, Glu) are not compatible with the enzyme’s function regardless of their size. For uncharged residues, size gains importance: Gln, Leu, Ile, Phe, Tyr and Trp mutants replicated poorly or not at all while Asn, Ser, Ala, Val and Pro mutants showed near WT replication levels, yet mutation to Gly was deleterious. Even more interestingly, positively charged residues support viral replication regardless of their size (Lys, Arg and His), in agreement with the observation that some polymerases such as that of SARS-CoV-2 carry a Lys residue at the corresponding position. Interestingly, the effect of a negative or bulky residue at position 109 could be mitigated by a second site, Asp106-Gly mutation (Fig. 5 and supplemental Table A1). Taken together these observations indicate Thr109 might play a role both in the structure and the net charge of the loop, likely guiding the RNA through the channel towards the catalytic site.

As phosphomimetic mutations strongly affect RdRp activity, the phosphorylation of either residue (Thr109-Ser110 / Ser502 for SARS-CoV-2) that can occur in infected cells would most likely inhibit viral replication. Although phosphorylation of Thr109/Ser110 was not prominent, it was detected in independent infection experiments and might occur in specific circumstances or in specific cell types.

These observations raise intriguing questions about the physiological function of 3D^pol^ phosphorylation: why are these phosphorylatable residues conserved when our results indicate they could be replaced by a non-phosphorylatable alanine without reducing the polymerase’s activity? Some hypotheses can be proposed:

i. Phosphorylation might occur preferentially in certain cell types where viral replication needs to go undetected by cellular sensors, thereby escaping immune defences, avoiding lytic replication, and allowing non-lytic viral transmission to other cells.
ii. Phosphorylation might serve as a switch between translation and replication. In ss(+)RNA viruses, the genome serves both as a platform for protein production and as a template for the production of negative sense RNA, both of which cannot take place at the same time on the same RNA molecule. The phosphorylation of the polymerase might intervene at a specific time of the viral cycle when translation needs to be favoured.
iii. Phosphorylation could also play a role in 3D^pol^ or 3CD protein turn-over, subcellular localization, partner(s) binding or activity. Our mutants could be low fidelity polymerases, which our assays were not designed to probe for. It has been observed at numerous occasions and in many different viruses that mutations on the fingers domain, sometimes far from the catalytic centre, can modify a polymerase’s fidelity, sometimes with the low fidelity mutant being faster but never more fit than the WT counterpart in competition assays [20, 21].

Yet, it is intriguing that polymerases might have evolved to have their activity silenced by cellular kinases.

It is also noteworthy that these residues are close to the highly conserved proline 114 (Fig. 2D) that is observed in a *cis* peptide bond conformation in picornaviral polymerases [22] and in many SARS-CoV-2 polymerase (PDB structures 9PYW, 7BW4 and 8SQK but not 7CTT). It has been postulated that a *cis-trans* isomerization of the proline would result in the protrusion of the Gly112–Gly119 segment (TMEV numbering) from the polymerase surface, and such a conformational change may be triggered by Ser110 phosphorylation or it may serve a regulatory function that limits kinase access to Ser110.

Our luciferase reporter assays done with SARS-CoV2 point towards a similar role between SARS-CoV2 nsp12 Lys501-Ser502 and TMEV 3D^pol^ Thr109-Ser110. Given the role of motif G in RdRps of (+)ssRNA viruses, it is reasonable to expect that polymerases from other viral families would react similarly. Polymerases being indispensable in the viral cycle of RNA viruses, our observations might serve the development of a large-spectrum antiviral non-nucleoside inhibitors (NNI) targeting the polymerase. In this sense, several NNIs have been developed that target the template entry channel. Our results, aside from showing that these residues are readily accessible (as they can undergo phosphorylation), could provide valuable information for drug design:

GPC-N114 was described as a NNI capable of inhibiting several enteroviruses and even EMCV, a member of the genus *Cardiovirus* [23]. The compound binds in a pocket at the bottom of the template entry channel, mimicking the position of the template acceptor nucleotide, which will base-pair with the incoming nucleotide. Interestingly, the analysis of its position in co-crystals with CVB3 3D^pol^ showed a direct interaction with Thr114, the TMEV Thr109 homolog [23].

NITD-1 was identified by high throughput screening as being able to selectively inhibit dengue virus (DENV) non-structural protein 5 (NS5, the RdRp). UV-crosslink followed by mass spectrometry analysis, docking and molecular simulation indicated the compound binds to the template entry channel of the polymerase at the intersection between the fingers and thumb [24].

NITD-640 binds to DENV1 NS5^pol^ motif F and is able to inhibit DENV1 NS5^pol^ *de novo* initiation and elongation. Its exact distance from motif G and specifically Ala417-Ala418 is impossible to determine due to the structure’s flexibility when NS5 is crystalized with NITD- 640. However, an alignment of the structure of DENV3 NS5 with the compound (PDB: 6XD1) with DENV3 NS5 without the compound (PDB: 5CCV) show that NITD-640 most likely displaces motif G when its bound to the polymerase [25].

Identification of phosphorylatable residues that impact polymerase activity might thus be valuable to optimize the design of molecules known to bind in the vicinity of the phospho- site to increase their inhibitory activity. These proteins being essential for the viral cycle of any RNA virus, our phosphorylation data might give rise to new antiviral strategies to lessen the burden of viral infections.

## Materials and methods

### Cell culture

293T cells [26] were kindly provided by Frédéric Tangy (Pasteur Institute, Paris, France). 293T, L929 (ECACC, 85011425), Neuro2A (ECACC, 89121404) cells were maintained in Dulbecco’s modified Eagle medium (DMEM) (BioWest) containing 4.5 g/L glucose, supplemented with 10% fetal calf serum (FCS) (Sigma). Raw264.7 cells (ATCC No. TIB-71) were kindly provided by Véronique Kruys (Université Libre de Bruxelles) and maintained in hydrophobic Petri dishes in DMEM containing 4.5 g/L glucose, supplemented with 5% Myoclone fetal calf serum (Gibco-BRL Myoclone Super Plus-Bovine Serum; catalog no. 10081-071) to limit macrophage activation [27] and 2 mM HEPES. Three days before infection, cells were cultured in media containing FCS instead of Myoclone serum; for the infections, cells were seeded in adherent dishes. BHK-21 cells (ATCC, CCL-10) and its derivate BHK-21-LgBiT were cultured in Glasgow’s minimum essential medium (GMEM) (Gibco) supplemented with 10% newborn calf serum (NCS) and 2.95 g/L tryptose phosphate broth.

All the cells mentioned above were cultured with 100 U/mL penicillin, 100 μg/mL streptomycine and maintained at 37 °C in 5% CO_2_. A549-hACE2-TMPRSS2 (InvivoGen, San Diego, CA, USA) were kept in DEMD (Gibco) supplemented with 10% FCS (HyClone^TM^), 0.5 µg/mL puromycin (InvivoGen) and 100 µg/mL hygromycin (InvivoGen).

### 293T cells transfection with pcDNA3 vectors coding for 3CD protein

293T cells (∼70% confluent) were transfected with pcDNA3 plasmids encoding the different 3CD mutants using Lipofectamine 2000 (Invitrogen Cat. #11668030). Cells were transfected following manufacturer’s instructions in 24-well plates with 3.5 µg of plasmid expressing 3CD for 24 h (see plasmids list in supplemental Table A2). They were lysed with 100 µL of Laemmli buffer and lysates were heated for 5 min at > 95 °C and stored at −20 °C.

### Western Blot

Proteins in Laemmli buffer were heated at >95 °C for 5 min before being run in 10% glycine SDS-PAGE. Proteins were then transferred to PVDF membranes. Membranes were first blocked with TBS-0.1% Tween 5% milk (Regilait) for 1 h at room temperature (RT) and then incubated over-night with primary antibodies at proper dilution in TBS-0.1% Tween 5% milk. The following primary antibodies were used: anti-3D^pol^ (produced in-house, see [28] 1/1,000) and anti-β actine (Sigma A5441 1/10,000). Membranes were washed three times with TBS-0.1% Tween 20 for 15 min before incubation with secondary antibodies in TBS-0.1% Tween 5% milk for 1 h at RT. Secondary antibodies used: HRP-conjugated anti-rabbit (Dako P0448 1/5,000), HRP-conjugated anti-mouse (Dako P0447,1/5,000). Membranes were washed three times with TBS-0.1% Tween 20, then once with TBS and developed with SuperSignal West chemiluminescence substrate (Pico or Dura, ThermoScientific). Images were acquired with a cooled CCD camera (Odyssey FC-LiCor).

### Production of BHK-21-LgBiT cells

pFW31 is a lentiviral vector constructed by inserting the nLuc LgBiT coding region from pFC34K (Promega) in pTM897 [12], a derivative of pCCLsin.cPPT.hPGK.GFP.pre [29]. pFW31 expresses nLuc LgBiT from a PGK promoter.

Lentiviruses were produced in 293T cells grown in a well of a 6-well plate by cotransfection of 0.75 μg of pMD2-VSV-G (VSV-glycoprotein), 1.125 μg of pMDLg/pRRE (Gag-Pol), 0.625 μg of pRSV-Rev (Rev), and 2.5 μg of pFW31, using the transfection reagent TransIT- LT1 (Mirus Bio). Supernatants containing the lentivirus were collected 48 h post transfection and filtered through a 0.45 μm filters.

For transduction, 10,000 BHK-21 cells grown in a well of a 24-well plate were infected twice with 150 μl of lentivirus. The transduced cells were cloned and 6 clones were tested by measuring their luciferase activity twelve hours after mock infection or after infection with virus MW2 (MOI 0.01 PFU/cell) expressing the nanoluciferase HiBiT. The clone with the highest infected/non-infected luciferase signal ratio was kept.

### TMEV viruses

For the mass spectrometry analysis, L929 cells were infected with TMEV KJ6, a variant of the persistent DA strain (DA1 molecular clone) adapted to grow on L929 cells [11].

Raw264.7 cells were infected with TMEV VV18, a variant of DA1 molecular clone carrying macrophage-adapted capsid mutations (VP2 G162S, VP1 V256E and VP1 S257Y)[12]. The Thr109 Arg (CAD8) mutant used in the competition assays was directly derived from the DA1 molecular clone. A list of the different plasmids, their mutations and name is provided in supplemental Table A2.

Viruses were produced from plasmid constructs containing the full-length viral cDNA sequences. To this end, BHK-21 cells were electroporated (1500 V, 25 μFd, no shunt resistance) with *in vitro* transcribed viral RNA (RiboMax transcription kit P1300, Promega). Supernatants were collected 48-72 h after electroporation, when cytopathic effect was complete. After 2 to 3 freeze-thaw cycles, the supernatants containing the virus were clarified by centrifugation at 1258x g for 20 min and viruses were stored at -80 °C. VV18 virus was further purified on a sucrose cushion as described in [12]. Final viral stocks (BHK supernatants or purified) were titrated by plaque assay in BHK-21 cells as described in [30].

### SARS-CoV-2 virus

SARS-CoV-2 VOC Alpha (lineage B.1.1.7; hCoV104 19/Belgium/rega-12211513/2020; EPI_ISL_791333) was grown on Calu-3 cells and harvested four days post-infection (dpi). Viral titers were determined using the Reed and Muench method and expressed as the 50% tissue culture infectious dose (TCID_50_) per ml. All research involving SARS-CoV-2 was conducted in the high-containment BSL3 facilities of the KU Leuven Rega Institute, under licenses AMV 30112018 SBB 219 2018 0892 and AMV 23102017 SBB 219 2017 0589, following institutional guidelines.

### Infections

Cells were seeded and grown to a confluency of 80%. Before infection, cells were rinsed with serum free medium (SF) and left for 1h in SF medium containing virus, after which at least 2 volumes of medium containing 10% of serum (SC medium) were added. For the competition assays, cells were rinsed once with SC medium, once with SF medium and left in medium containing 1% of serum.

Once the time of infection elapsed, cells were lysed either with IP lysis solution (see below), with Laemmli buffer (if crude lysates were used), or with a solution compatible with downstream RNA extraction.

### Immunoprecipitations

Infected cells were rinsed with ice-cold PBS and lysed for 15 min on ice with lysis buffer (for 3D^pol^ IP: Tris pH 8 50 mM, NaCl 150 mM, NP40 1%, DTT 0.25 mM, Sodium deoxycholate 0.2%; for nsp12 IP: Tris pH 8 50 mM, NP40 0.1%, SDS 0.1%) supplemented with 0.3 mg/mL of RNAse A (Roth 756) and protease and phosphatase inhibitor (ThermoScientific A32961, one tablet in 3.3 mL of lysis buffer). The lysates were homogenized by 10 passages through 21G needles and centrifuged at 12,000xg for 10 min at 4 °C. Pellets were sometimes collected for mass spectrometry analysis in which case, after cleared lysates were recovered, pellets were resuspended in Laemmli buffer. Lysates were then diluted 3 times with dilution buffer (Tris pH 8 50 mM, NaCl 20 mM, EDTA 0.2 mM, Glycerol 5%), pre-cleared by incubation with proteinA/G coupled magnetic beads (Pierce 88803) for 30 min at 4 °C, before being incubated for 2 hours with A/G magnetic beads coupled to either nsp12 or 3D^pol^ specific antibodies. After incubation, beads were rinsed 3 times with diluted lysis buffer, resuspended in Laemmli buffer (regular Laemmli was used for nsp12 IPs while 3x concentrated Laemmli buffer with no reducing agent was used for 3D^pol^) and heated (5 min at >95 °C for nsp12, 15 min at 37 °C for 3D^pol^) to allow proteins elution from the beads. Supernatants were conserved at −20 °C.

Crude cells lysates (Laemmli lysed), IP products or pellets were heated at >95 °C for 5 min and run on 8% (nsp12) or 10% (3D^pol^) Tris-glycine SDS-PAGE. Gels were stained with Coomassie blue (Thermo scientific ref. 24620) and gel fragments corresponding to the expected sizes for 3D^pol^ or nsp12 (55 kDa or 100 kDa respectively) were cut out based on protein ladders.

### Antibodies

Home-made anti-3D^pol^ and anti-nsp12 antibodies were obtained by injection of recombinant 3D^pol^ or nsp12 proteins into rabbits. Antibodies were then affinity-purified. Details of this procedure have been described in [28].

Recombinant nsp12 was obtained from His-tagged nsp12 protein purified on HisTrap FF column (Cytiva) as described elsewhere [31].

### Mass Spectrometry

Proteins were separated in a 10% acrylamide gel. Protein bands were visualized by colloidal Coomassie Blue staining and in-gel digested with trypsin. Peptides were extracted with 0.1% TFA in 65% ACN and dried in a speedvac.

Peptides were dissolved in solvent A (0.1% TFA in 2% ACN), directly loaded onto reversed-phase pre-column (Acclaim PepMap 100, Thermo Scientific) and eluted in backflush mode. Peptide separation was performed using a reversed-phase analytical column (Acclaim PepMap RSLC, 0.075 x 250 mm,Thermo Scientific) with a linear gradient of 4%-27.5% solvent B (0.1% FA in 98% ACN) for 40 min, 27.5%-50% solvent B for 20 min, 50%-95% solvent B for 10 min and holding at 95% for the last 10 min at a constant flow rate of 300 nl/min on an Vanquish Neo system. The peptides were analyzed by an Orbitrap Fusion Lumos tribrid (ThermoFisher Scientific). The peptides were subjected to NSI source followed by tandem mass spectrometry (MS/MS) in Fusion Lumos coupled online to the nano-LC. Intact peptides were detected in the Orbitrap at a resolution of 120,000. Peptides were selected for MS/MS using HCD setting at 30, ion fragments were detected in the linear ion trap (LUMOS). A data-dependent procedure that alternated between one MS scan followed by MS/MS scans was applied for 3 seconds for ions above a threshold ion count of 2.0E4 in the MS survey scan with 40.0s dynamic exclusion. The electrospray voltage applied was 2.1 kV. MS1 spectra were obtained with an AGC target of 4E5 ions and a maximum injection time of 50 ms, and MS2 spectra were acquired with an AGC target of 5E4 ions and a maximum injection time set to dynamic. For MS scans, the m/z scan range was 375 to 1800. The resulting MS/MS data was processed using Sequest HT search engine within Proteome Discoverer 2.5 SP1 against a *Homo sapiens* or *Mus musculus* protein database obtained from Uniprot and containing the TMEV or SARS-CoV2 sequence. Trypsin was specified as cleavage enzyme allowing up to 2 missed cleavages, 4 modifications per peptide and up to 5 charges. Mass error was set to 10 ppm for precursor ions and 0.1 Da for-fragment ions.

Oxidation on Met (+15.995 Da), conversion of Gln (-17.027 Da) or Glu (- 18.011 Da) to pyro-Glu at the peptide N-term and phosphorylation on Ser, Thr, Tyr (+79.966 Da) were considered as variable modifications. False discovery rate (FDR) was assessed using Percolator and thresholds for protein, peptide and modification site were specified at 1%. For abundance comparison, abundance ratios were calculated by Label Free Quantification (LFQ) of the precursor intensities within Proteome Discoverer 2.5 SP1.

The mass spectrometry proteomics data have been deposited to the ProteomeXchange Consortium via the PRIDE [32–34] partner repository with the dataset identifier PXD071274 (doi: DOI: 10.6019/PXD071274).

### Isolation and sequencing of revertant viruses

BHK-21 cells were electroporated (1,500 V, 25 μFd, no shunt resistance) with *in vitro* transcribed viral RNA (RiboMax transcription kit P1300, Promega). Electroporated BHK-21 cells were seeded in 75 cm^2^ dishes containing BHK-21 cells at 1/6 confluency. Six hours post seeding, cells were covered with 0.5% agarose in DMEM and left for 48 h at 37 °C, 5% CO_2_. Plaques were pipetted out from the agarose, suspended in 300 µL of SF-DMEM for at least 6h at 4 °C, and used to infect fresh BHK-21 cells for 3 days. RNA was recovered as described in [35] and reverse-transcribed, PCR amplified using TM16 (ACC CGA CGT GGT TGG TGT GG) and TM272 (GTC AGT TCC AAT TGC AGA TCC) and the PCR product was purified (Promega A19282) before being Sanger-sequenced (Eurofins) using primer TM3242 (TGG TCT GGA ATT TGG AGG CA). All DNA sequence analyses and assemblies were performed in ApE [36].

### Competition assays

Cells were seeded in 24-well plates and infected with a 1:1 mixture of CAD8 and DA1 viruses in a volume of 200 μL. The initial MOI depended on the cell type infected (Neuro 2A, BHK-21, 1 PFU/cell, L929: 0.1 PFU/cell, RAW264.7: 5 PFU/cell). After infection, cells were left in 500 µL of culture medium until CPE occurred. 400 µL of supernatant were collected, cleared with a 20 min centrifugation and kept at -80 °C or used in part for the next round of infection; the leftover (medium + cells) was recovered for downstream RNA extraction. For the following infections, the same protocol was followed but diluted supernatants were used instead of a viral mixture. The dilution factor of the supernatants depended on the cell type, a ½ dilution meaning cells were infected with 100 µL of supernatant supplemented with 100 µL of SF medium, the dilutions were as follow: ½ for RAW264.7 cells, ¼ for Neuro2A cells and 1/10 for BHK-21 and L929 cells.

Isolated RNA was reverse-transcribed and PCR amplified with TM16 and TM272 primers. The PCR products were digested with BglII (Fermentas ER0082) for 3 hours at 37 °C, loaded in Tris/Borate/EDTA (TBE), 2% agarose gels containing ROTI gel stain (Carl Roth ref. 3865) and migrated for 1 h at 90 V. The DNA fragments were visualized using a Proxima imaging platform from Isogen.

### Luciferase assays

For the TMEV luciferase assays, 600,000 BHK-21-LgBiT cells, washed and suspended in 700 μl of PBS, were electroporated (1500 V, 25 μFd, no shunt resistance) with *in vitro* transcribed viral RNA (Promega P207E). Electroporated BHK-21 cells were diluted 4 times in GMEM supplemented with 10% NCS and 2.95 g/L tryptose phosphate broth and seeded in 96 well plates (40,000 cells per well or 1/15^th^ of the total).

For the SARS-CoV-2 luciferase assays, the BACs containing the cDNA of SARS-CoV-2 were diluted in pOPTG plasmid DNA (ratio 20:80 SARS-CoV-2:pOPTG) before transfection. 293T cells (40,000 per well) were transfected with 100 ng of DNA mixture using Lipofectamine 2000 (Thermofisher scientific). DNA-Lipid complexes were produced following the manufacturers’ instructions, cells were added to the complexes and left 5 min at RT before being seeded in 96 wells plates. Actinomycin D (ROTH 8969) diluted in DMSO was added directly to the wells at a final concentration of 0.5 μg/mL.

For each luciferase measurement, cells were lysed with the promega kit (P1130) following the manufacturer’s instructions. Luciferase signal was measured using a GloMax luminometer (Promega).

## Statistical analysis

Statistical analysis was performed as indicated using Prism 10 (GraphPad Software). Error bars represent the standard deviation. *P* values: \*\*\**P* < 0.001, \*\**P* < 0.01, \**P* < 0.05, ns = not significant.

## Protein sequence alignments

Sequences of representative Cardiovirus isolates were obtained from the international committee of taxonomy of viruses (ICTV) website (https://ictv.global/report/chapter/picornaviridae/picornaviridae/cardiovirus). Picornaviruses 3D^pol^ sequences were obtained from public sequence datasets available on the GenBank platform using the Basic Local Alignment Search Tool (BLAST) [37] developed by the NIH. The search was done using TMEV sequence as a query for refseq_reference_genomes within taxid 12058.

Sequences were aligned using Clustal Omega Multiple Sequence Alignment (MSA) [38] and visualised using Jalview 2.11.4.1 [39] or WebLogo [40].

## Structure prediction and visualization

Structures were predicted using AF2 via ColabFold [41] and visualized using PyMol [42].

## Data availability

All data are contained in the article. Biological material is available upon reasonable request.

## Conflict of interest

The authors declare that the research was conducted in the absence of any commercial or financial relationships that could be construed as a potential conflict of interest.

## Acknowledgement

We are very grateful to Prof. Andrès Merits (Institute of Technology, University of Tartu, Estonia) for the gift of the BAC pCCI-4K-SARS-CoV-2-NLuc, which allowed us to derive the replicon-encoding BAC. We also thank Frédéric Sorgeloos for his help in data analysis and insightful ideas, Tina van Buyten and Elke Maas for the maintenance of the cell lines at BSL3, and Mathilde Wieme for the construction of the Thr109 and Ser110 reporter viruses.

## Funding statement

CD and BLP were the recipient of a fellowship from the FNRS-FRIA (National fund for Scientific Research) and were then supported by the EOS (IDs: 30981113 and 40007527); Research was supported by FNRS (PDR T.0154.23), EOS (EOS ID: 40007527), and Loterie Nationale through support to the de Duve Institute. The funders had no role in study design, data collection and analysis, decision to publish, or preparation of the manuscript.

## Authors contribution

Conceptualization: C.D., T.M., B.L.-P. Methodology: C.D., T.M., B.L.-P. Validation: D.V., C.D., T.M. Formal analysis: C.D., D.V. Investigation: C.D., BL-P, O.B.P., Y.A.A., G.H. Resources, Project administration and Funding acquisition: T.M., O.B.P., Y.A.A. Supervision: T.M. Data Curation: C.D., T.M., D.V., B.L.-P. Visualization: C.D. Writing-Original Draft and Visualization: C.D. Writing: C.D., T.M. Review & Editing: C.D., Y.A.A., O.B.P., B.L.-P., T.M.

